# A Mixed Periodic Paralysis & Myotonia Mutant, P1158S, Imparts pH Sensitivity in Skeletal Muscle Voltage-gated Sodium Channels

**DOI:** 10.1101/164988

**Authors:** Mohammad-Reza Ghovanloo, Mena Abdelsayed, Colin H. Peters, Peter C. Ruben

## Abstract

Skeletal muscle channelopathies, many of which are inherited as autosomal dominant mutations, include both myotonia and periodic paralysis. Myotonia is defined by a delayed relaxation after muscular contraction, whereas periodic paralysis is defined by episodic attacks of weakness. One sub-type of periodic paralysis, known as hypokalemic periodic paralysis (hypoPP), is associated with low potassium levels. Interestingly, the P1158S missense mutant, located in the third domain S4-S5 linker of the ‘‘skeletal muscle’’ voltage-gated sodium channel, Nav1.4, has been implicated in causing both myotonia and hypoPP. A common trigger for these conditions is physical activity. We previously reported that Nav1.4 is relatively insensitive to changes in extracellular pH compared to Nav1.2 and Nav1.5. Given that intense exercise is often accompanied by blood acidosis, we decided to test whether changes in pH would push gating in P1158S towards either phenotype. Our results indicate that, unlike in WT Nav1.4, low pH depolarizes the voltage-dependence of activation and steady-state fast inactivation, decreases current density, and increases late currents in P1185S. Thus, P1185S turns the normally pH-insensitive Nav1.4 into a proton-sensitive channel. Using action potential modeling we also predict a pH-to-phenotype correlation in patients with P1158S. We conclude that activities which alter blood pH may trigger myotonia or periodic paralysis in P1158S patients.

**SIGNIFICANCE STATEMENT:** Voltage-gated sodium channels (Nav) contribute to the physiology and pathophysiology of electrical signaling in excitable cells. Nav subtypes are expressed in a tissue-specific manner, thus they respond differently to physiological modulators. For instance, the cardiac subtype, Nav1.5, can be modified by changes in blood pH; however, the skeletal muscle subtype, Nav1.4, is mostly pH-insensitive. Nav1.4 mutants can mostly cause either hyper-or hypo-excitability in skeletal muscles, leading to conditions such as myotonia or periodic paralysis. P1158S uniquely causes both phenotypes. This study investigates pH-sensitivity in P1158S, and describes how physiological pH changes can push P1158S to cause myotonia and periodic paralysis.

## INTRODUCTION

Electrical signaling is an integral part of many processes including the heart beat and voluntary movement. This kind of signaling depends on a group of transmembrane proteins called voltage-gated ion channels (1). Voltage-gated sodium channels are responsible for the initiation and propagation of action potentials in most excitable cells. These channels are hetero-multimeric proteins composed of a large voltage-sensing and pore-forming α-subunit, and smaller β-subunits (2). The α-subunit family is composed of 9 subtypes, which are expressed in different parts of the body (3). Nav1.4, highly expressed in skeletal muscle, is encoded by the *SCN4A* gene. Mutations in *SCN4A* often lead to either gain-of-function or loss-of-function phenotypes (4).

Although most Nav1.4-mutants depolarize the sarcolemma membrane, this depolarization can result in either hyper-or hypo-excitability (5). The manifestation of Nav-channelopathies occurs through changes in membrane excitability, and these changes underlie various clinical syndromes. Traditionally, muscular channelopathies are classified as either non-dystrophic myotonias or periodic paralysis (6, 7). Most of these channelopathies are either sporadic *de novo* or inherited autosomal dominant mutations (8).

The phenotypic expression of most muscle channelopathies takes place within the first two decades of life, and many afflicted individuals suffer from life-long symptoms such as muscle stiffness, weakness, or pain (9). Despite their high penetrance, these mutations can show clinical variability within different familial groups or even a single family (9).

Most gain-of-function mutations in *SCN4A* result in myotonic syndromes. Myotonia is defined by a delayed relaxation after muscle contraction (9, 10). In myotonia, there is an increase in muscle membrane excitability in which even a brief voluntary contraction can lead to a series of action potentials that can persist for several seconds following the termination of motor neuron activity. This phenomenon is perceived as muscle stiffness (9). The global prevalence of non-dystrophic myotonias is estimated at 1: 100,000 (11). Although this condition is not considered lethal, it can be extremely life-limiting due to the multitude of muscle contractility problems it can cause. Given the characteristic hyper-excitability of myotonia, most therapeutics are targeted against membrane excitability (12–14).

Until recently, loss-of-function Nav1.4-mutants were believed to cause periodic paralysis. This was due to most periodic paralysis-causing mutations neutralizing positively-charged residues in voltage sensors. Newer studies described a cation leak with characteristics similar to the ω-current in *Shaker* potassium channels. These findings indicate a form of gain-of-function resulting in periodic paralysis (15–18). Some channelopathies are triggered by low serum potassium levels and manifest in episodes of extreme muscle weakness. These are known as hypokalemic periodic paralyses (hypoPP) (19). The onset of hypoPP occurs between the ages of 15 and 35, and the prevalence of hypoPP is also estimated to be 1: 100,000. HypoPP can be triggered by external factors such as stress, high-sugar diet, and exercise (19). A serum potassium concentration less than 3 mM may trigger hypoPP (20).

P1158S in Nav1.4 is a missense mutant in the S4-S5 linker of domain (D) III that is the result of a C to T mutation at position 3549 in *SCN4A* (21). P1158S is described to cause both myotonia and hypoPP. The ability to cause two seemingly contrasting syndromes is shared between P1158S and some mutants of the cardiac sodium channel, Nav1.5, which cause both the Brugada and long QT syndromes (22). Our group recently showed that physiological triggers may be important in modulating disease phenotype in the E1784K mutant in Nav1.5 (23), leading us to hypothesize a possible role of similar triggers in P1158S.

Exercise is a common external trigger for both myotonia and hypoPP (9, 10, 19). Physical activity results in delayed relaxation in myotonic patients and extreme weakness in hypoPP patients. Exercise is normally accompanied by changes in extracellular pH, which can modulate sodium channel function (23–25). We therefore hypothesized that pH alterations may push P1158S towards either myotonia or hypoPP.

Changes in extracellular pH levels modulate the activities of Nav1.2 and Nav1.5; however, Nav1.4 is relatively pH-insensitive (25). One theory behind the pH-sensitivity of Nav1.5 is the presence of a cysteine (C373) residue on the outer vestibule of DI (24, 26). The increase in H^+^ concentration results in the protonation of this cysteine, which creates a positive charge outside the pore. This positive charge causes an electrostatic repulsion, blocking sodium ions from going through the pore. This cysteine is missing in Nav1.4 (26). However, there are other residues on the extracellular side of Nav1.4 that may get protonated (26, 27). We postulated that one reason for the apparent pH-insensitivity of Nav1.4 relative to other sodium channels might be due to hydrophobic amino acids protecting the protonate-able functional groups from becoming protonated at low pH.

The serine functional group is floppier than the highly-restrained proline functional group. Furthermore, mutational studies suggest that in Nav1.4, DIII activates before DI and DII (28). Given the location of P1158S on the DIII S4-S5 linker, along with the nature of the mutation, and because DIII activates first, we predicted that the increased mobility of DIII in P1158S might expose residues protected in WT Nav1.4 to protonation. Here we report the results of homology modeling, patch-clamp experiments, and action potential modeling. These results support our hypothesis and show P1158S imparts pH-sensitivity to Nav1.4 that underlies the dual myotonia/hypoPP phenotype.

## MATERIALS AND METHODS

### Cell Culture

Chinese Hamster Ovary (CHO) cells were transiently co-transfected with cDNA encoding eGFP and the β1-subunit and either wild-type Nav1.4 or P1158S α-subunit. Transfection was done according to the PolyFect transfection protocol. After each set of transfections, a minimum of 8-hour incubation was allowed before plating on sterile coverslips.

### Homology Modeling

Homology modeling was performed using the Swiss-Model server (29). The sodium channel structure developed by Ahuja et al., (2015) was used as a template against which the Nav1.4 sequence with and without P1158S were modeled (29, 30). Modeling was done according to the protocol established by Bordoli et al., (2008). PyMOL-pdb viewer was used for optimization and structure visualization.

### Electrophysiology

Whole-cell patch clamp recordings were performed in an extracellular solution containing (in mM): 140 NaCl, 4 KCl, 2 CaCl_2_, 1 MgCl_2_, 10 HEPES or MES (pH6.4). Solutions were adjusted to pH (6.4, 7.0, 7.4, 8.0) with CsOH. Pipettes were filled with intracellular solution, containing (in mM): 120 CsF, 20 CsCl, 10 NaCl, 10 HEPES or MES (pH6.4). All recordings were made using an EPC-9 patch-clamp amplifier (HEKA Elektronik, Lambrecht, Germany) digitized at 20 kHz via an ITC-16 interface (Instrutech, Great Neck, NY, USA). Voltage-clamping and data acquisition were controlled using PatchMaster/FitMaster software (HEKA Elektronik, Lambrecht, Germany) running on an Apple iMac. Current was low-pass-filtered at 10 kHz. Leak subtraction was performed automatically by software using a P/4 procedure following the test pulse. Gigaohm seals were allowed to stabilize in the on-cell configuration for 1 min prior to establishing the whole-cell configuration. Series resistance was less than 5 MΩ for all recordings. Series resistance compensation up to 80% was used when necessary. All data were acquired at least 1 min after attaining the whole-cell configuration. Before each protocol, the membrane potential was hyperpolarized to –130 mV to ensure complete removal of both fast inactivation and slow inactivation. All experiments were conducted at 22 °C.

### Activation Protocols

To determine the voltage-dependence of activation, we measured the peak current amplitude at test pulse potentials ranging from –100 mV to +80 mV in increments of +10 mV for 20 ms. Channel conductance (G) was calculated from peak I_Na_:

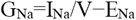

where G_Na_ is conductance, I_Na_ is peak sodium current in response to the command potential V, and E_Na_ is the Nernst equilibrium potential. Calculated values for conductance were fit with the Boltzmann equation:

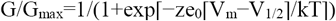

where G/G_max_ is normalized conductance amplitude, V_m_ is the command potential, z is the apparent valence, e_0_ is the elementary charge, V_1/2_ is the midpoint voltage, k is the Boltzmann constant, and T is temperature in °K.

### Steady-State Fast Inactivation Protocols

The voltage-dependence of fast inactivation was measured by preconditioning the channels to a hyperpolarizing potential of −130 mV and then eliciting pre-pulse potentials that ranged from −170 to +10 mV in increments of 10 mV for 500 ms, followed by a 10 ms test pulse during which the voltage was stepped to 0 mV. Normalized current amplitudes from the test pulse were fit as a function of voltage using the Boltzmann equation:

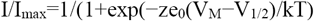

where *I*_max_ is the maximum test pulse current amplitude.

### Late Current Protocols

Late current was measured between 45 and 50 ms during a 200 ms depolarizing pulse to 0 mV from a holding potential of-130 mV. Pulses were averaged to increase signal-to-noise ratio. **Use-Dependent Inactivation Protocols**

Cells were held at −60 mV and repetitively stimulated with 2000, 20 ms test pulses to 0 mV. Peak currents from test pulses were normalized to the amplitude of the first current in the series and values were plotted versus the time of each pulse in seconds.

### Immunocytochemistry

CHO cells co-transfected with the β1-subunit and wild-type Nav1.4 or P1158S α-subunits were incubated for 24 hours. The transfections were done on glass coverslips. Non-transfected cells were also incubated, and were used as negative control. After this incubation period, cells were fixed in 4% paraformaldehyde for 10 minutes while being incubated on a shaker. Then, cells were incubated in 0.1% Triton X-100 for another 10 minutes. Cells were blocked in 10% goat serum for 30 to 45 minutes. Following this step, the cells were kept on Parafilm on top of a wet filter paper. Diluted primary anti-Nav1.4 (Rabbit polyclonal IgG, commercially available via Alomone) antibodies were added. The next day, cells were incubated in PBS for 10 minutes. Lastly, secondary IgG2a was added. Stained cells were studied using a Zeiss confocal microscope.

### Action Potential Modeling

Skeletal action potential modeling was based on a model developed by Cannon el al., (1993). All action potentials were programmed and run using Python. The modified parameters were based on electrophysiological results obtained from whole-cell patch-clamp experiments (31). The model accounted for activation voltage-dependence, steady-state fast inactivation voltage-dependence, and late sodium currents. The WT pH7.4 model uses the original parameters from the model. As the WT Nav1.4 was not sensitive to changes in pH, the WT models are identical for all pH values. P1158S models were programmed by shifting the midpoints of activation and fast inactivation from the original Cannon model by the difference between the values in P1158S experiments at a given pH and the average value in WT at all pH values. As persistent current was increased by the P1158S mutant, but was not significantly impacted by changes in pH, the simulated P1158S persistent current was the same at all pH values.

### Statistics

A two-factor analysis of variance (ANOVA) was used to compare the mean responses [activation, current density, steady-state fast inactivation, late currents, fast inactivation kinetics, use-dependent inactivation] between the channel variants. Channel variant had two levels (WT, P1158S) and pH had four levels (pH 6.4, 7.0, 7.4, 8.0). A one-factor analysis of variance (ANOVA) was used to compare the means membrane fluorescence intensities of the two channels and a negative control. *Post hoc* tests using the Tukey Kramer adjustment compared the mean responses between channel variants across conditions. A level of significance α = 0.05 was used in all overall *post hoc* tests, and effects with p-values less than 0.05 were considered to be statistically significant. All values are reported as means ± standard error of means for n cells.

### Analysis

Analysis and graphing were done using FitMaster software (HEKA Elektronik) and Igor Pro (Wavemetrics, LakeOswego, OR, USA). All data acquisition and analysis programs were run on an Apple iMac (Apple Computer). Statistical analysis was performed in JMP version 13. Confocal images were analyzed in ImageJ.

## RESULTS

### P1158S Alters the Hinge Angle of the DIII S4-S5 Linker

In the sliding-helix model of voltage-sensor movement during activation, when the potential across the cell membrane becomes depolarized, the voltage-sensor domain (VSD) undergoes an outward movement. This movement pulls on the S4-S5 linker and drags open the pore-domain (PD), leading to an influx of sodium ions (32). Based on this model we sought to determine whether P1158S alters the S4-S5 topology. We predicted that any topological change might alter the mobility of this linker and impart pH-sensitivity during acidosis.

To test our hypothesis, we performed homology modeling using the Swiss-Model server, accessible via the ExPASy web server, to model the sequence of both WT and P1158S against the Nav1.7 structure (Ahuja et al., 2015) (33). The Nav1.7 structure was used as a skeleton on which the WT and mutant-Nav1.4 sequences were placed (29). Our results show the critical position of P1158 in Nav1.4 on the hinge of DIII S4-S5 linker (**Fig. 1A-C**).

**Fig. 1.**
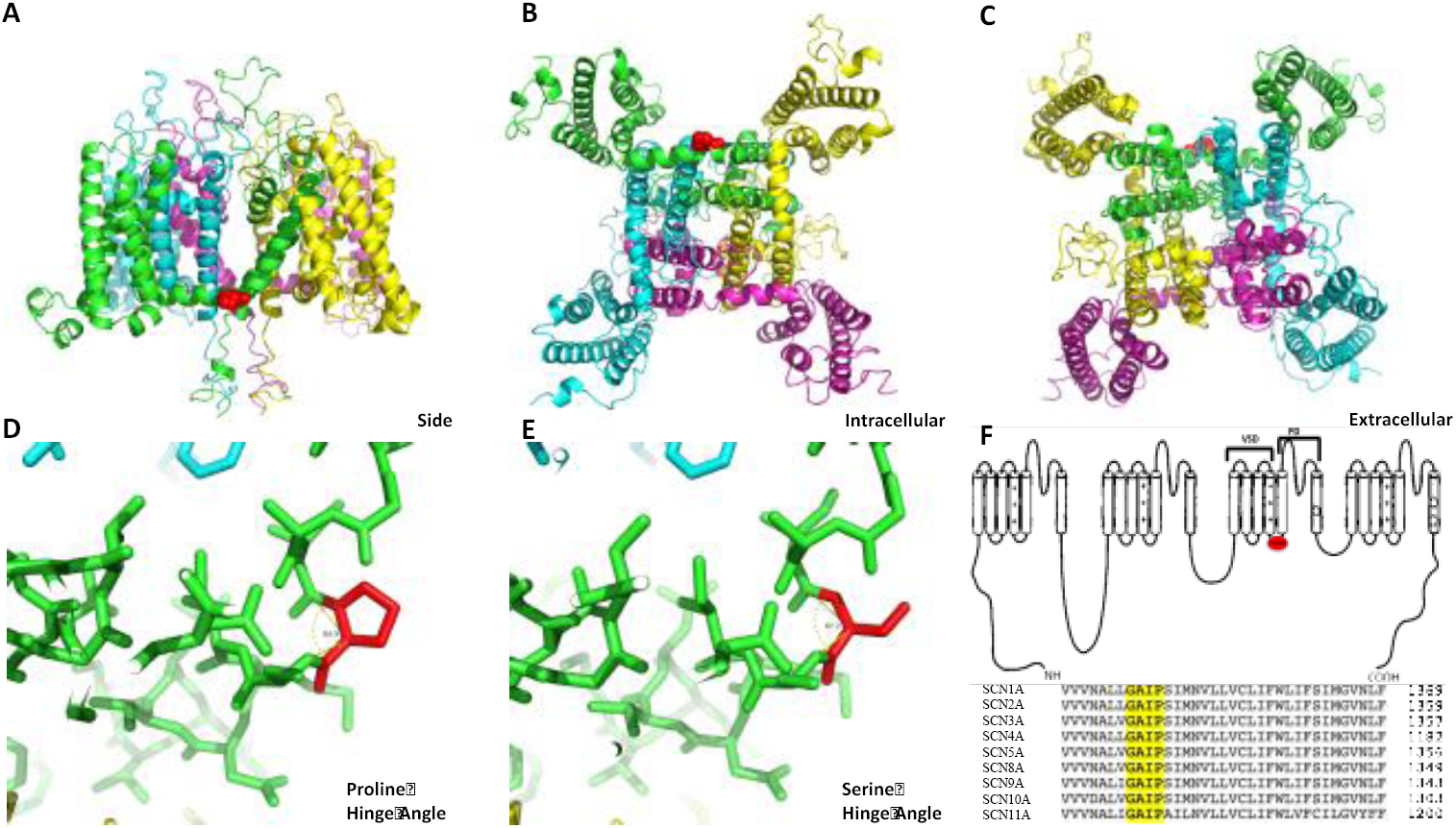
Homology model of sodium channel. **(A-C)** Show the views of the location of P1158 from the side, intracellular, and extracellular sides. The structure is colored by domain. P1158 is highlighted in red. **(D-E)** Show the hinge angle difference between having proline or serine at position 1158. Proline gives the hinge an 84.9° angle and serine gives the hinge an 87.2° angle. **(F)** Show the location of the P1158S mutation on a generic sodium channel schematic (highlighted in red), and the conservation of this part of Nav1.4, aligned with the rest of sodium channels.

Gain-and loss-of-function mutations may alter the atomic structure of sodium channels by causing connectivity rearrangements and changing interatomic bonds and topological angles (34). Therefore, we ran the model with the P1158S mutant to test whether there was an effect on the hinge angle of the S4-S5 linker. The hinge angle with proline was 84.9°; however, this angle was increased to 87.2° with the serine mutation (**Fig. 1D-E**). This suggests there is less angle strain on the S4-S5 linker when serine is present as opposed to proline. P1158 is conserved across all Nav subtypes, further suggesting its structural importance (**Fig. 1F**).

### P1158S Makes Nav1.4 Sensitive to Changes in Extracellular, but not Intracellular pH

We conducted whole-cell patch-clamp experiments to investigate whether the P1158S mutant in the S4-S5 linker increases the sensitivity of Nav1.4 to protons. We examined the effects of pH changes on channel activation in WT and P1158S by measuring peak channel conductance at membrane potentials between −100 and +80 mV (**Fig. 2H**). Neither decreasing nor increasing extracellular pH causes significant effects on the midpoint (V_1/2_) or apparent valence (z) of activation in WT channels (p > 0.05). In the P1158S mutant, however, there are significant shifts in the depolarized direction in the V_1/2_ of activation when extracellular pH is changed from pH8.0 to 6.4 (p = 0.0011) (**Fig. 2A-E**). We fitted the conductance V_1/2_ of P1158S using the Hill equation and WT using a flat line (**Fig. 2G**) (**Table S1**). The P1158S mutant has a pKa of 7.27 for the V_1/2_ of activation.

**Fig. 2.**
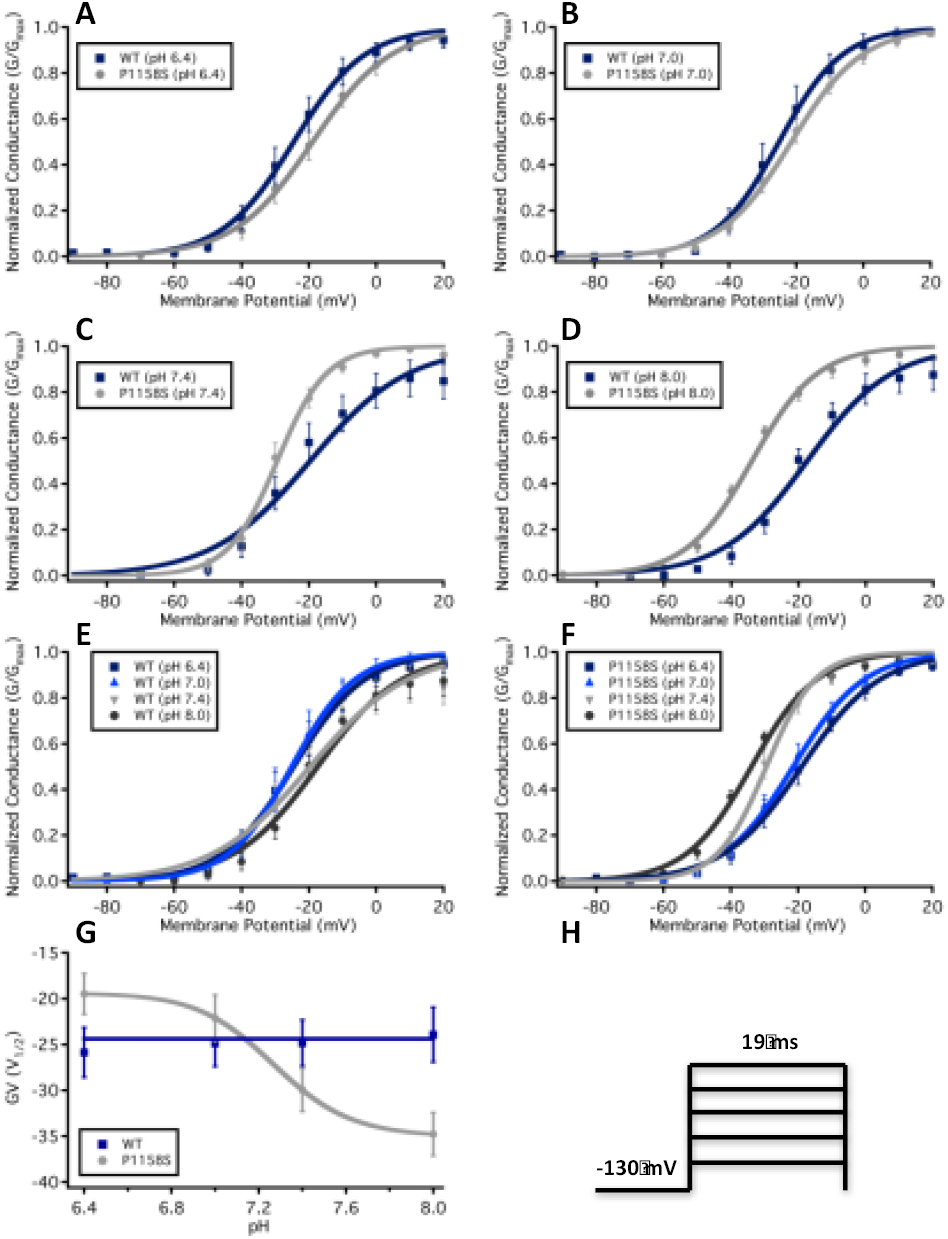
Normalized conductance plotted against membrane potential. **(A-D)** Show overlaps of WT (blue squares) and P1158S (grey circles) conductance at pH6.4 **(A)**, 7.0 **(B)**, 7.4 **(C)**, and 8.0 **(D)**. **(E-F)** Show conductance of each WT **(E)** and P1158S **(F)**. **(G)** Shows the voltage-dependence of activation as a function of pH and fitted with Hill equation (P1158S) or a straight line (WT). **(H)** Shows pulse protocols used to measure sodium channel conductance.

We measured current density from the ratio of peak current amplitude to the cell membrane capacitance (pA/pF). Representative current traces are shown in (**Fig. 3A-H**). Whereas the current densities of WT channels were not significantly affected by changes to extracellular pH (p > 0.05), the P1158S current density was significantly decreased at low extracellular pH (p = 0.0014) (**Fig. 3I**) (**Table S2**).

**Fig. 3.**
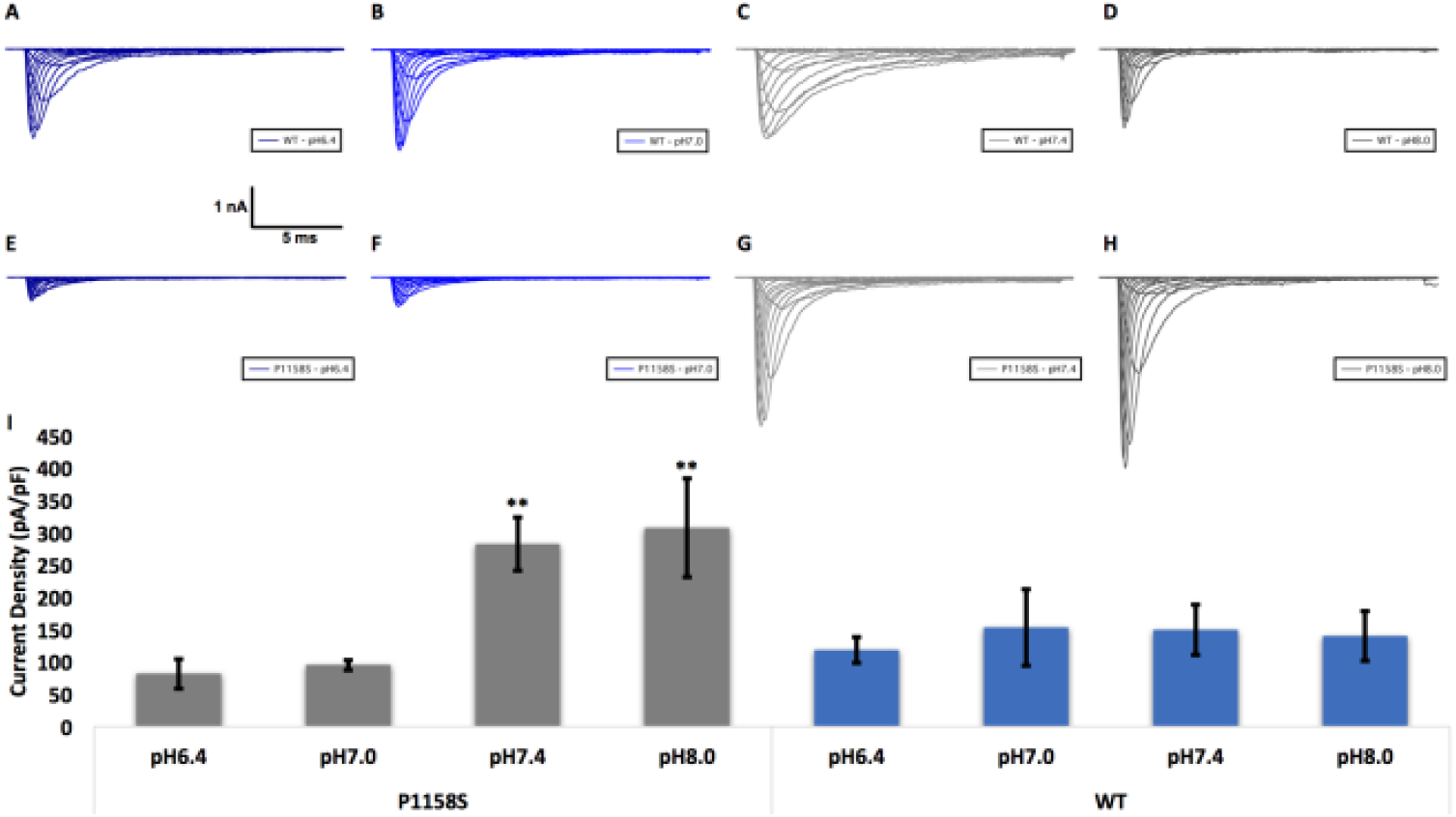
Current density measured in pA/pF. **(A-H)** Sample macroscopic sodium currents elicited by depolarizations between-100 and +80 mV. **(I)** Average current density of P1158S and WT-Nav1.4 at extracellular pH between 6.4 and 8.0. ** indicate p < 0.01.

After activation, the DIII-DIV linker, known to be the fast inactivation gate, binds to the inside of the channel within milliseconds and blocks the flow of sodium through the pore (35). We measured the voltage-dependence of fast inactivation using a standard pre-pulse voltage protocol (**Fig. 4H**). Normalized current amplitudes were plotted as a function of pre-pulse voltage (**Fig. 4A-F**). Like activation, the voltage-dependence of steady-state fast inactivation was depolarized by low extracellular pH in P1158S (p = 0.0027). Also, similar to activation, the WT V_1/2_ of fast inactivation was not significantly affected by changes in extracellular pH (p > 0.05) (**Table S3**). We fit fast inactivation V_1/2_ as a function of pH with either a Hill curve (P1158S) or a flat line (WT) (**Fig. 4G**). The P1158S V_1/2_ of fast inactivation has a pKa of 7.63. We also measured the open-state fast inactivation time constant, which did not differ significantly between channels or pH points (p > 0.05) (**Fig. S1**) (**Table S4**).

**Fig. 4.**
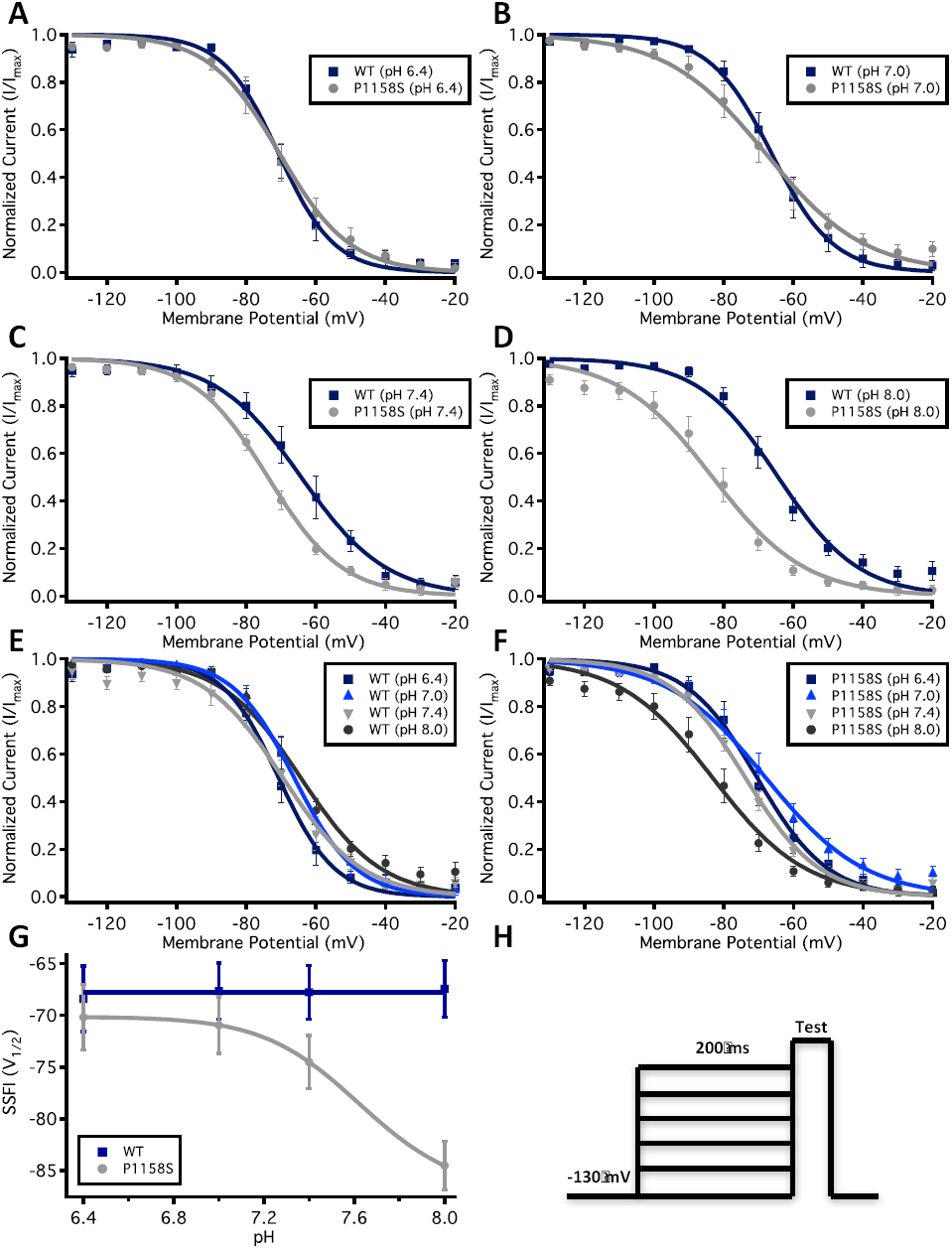
Voltage-dependence of steady-state fast inactivation as normalized current plotted against membrane potential. **(A-D)** Show the voltage-dependence of fast inactivation of WT (blue squares) and P1158S (grey circles) at pH6.4 **(A)**, 7.0 **(B)**, 7.4 **(C)**, and 8.0 **(D)**. **(E-F)** Show voltage-dependence of fast inactivation of each WT **(E)** and P1158S **(F)**. **(G)** Shows the voltage-dependence of steady-state fast inactivation as a function of pH and fitted with a Hill equation (P1158S) or a straight line (WT). **(H)** Shows voltage protocols used to measure the voltage-dependence of steady-state fast inactivation.

Total sodium current can be divided into two components: peak (INaP) and late currents (INaL). Whereas INaP refers to the maximum amount of sodium ions going through the channels during the open state, the smaller INaL is a manifestation of destabilized fast inactivation. We show representative normalized current traces for both channels across all conditions (**Fig. 5A-D**). We measured the percentage of INaL by dividing the maximum INaL between 145 ms and 150 ms ranges by INaP. Our results indicated that WT INaL is not significantly sensitive to changes in pH (p > 0.05). In contrast, P1158S showed a channel effect with a greater fraction of late current compared to WT (p = 0.0041) (**Fig. 5E**) (**Table S5**). This is an important result because previous studies suggested that late sodium current may be the basis of repetitive action potential firing and, consequently, myotonia in skeletal muscle fibers (5, 31, 36, 37).

**Fig. 5.**
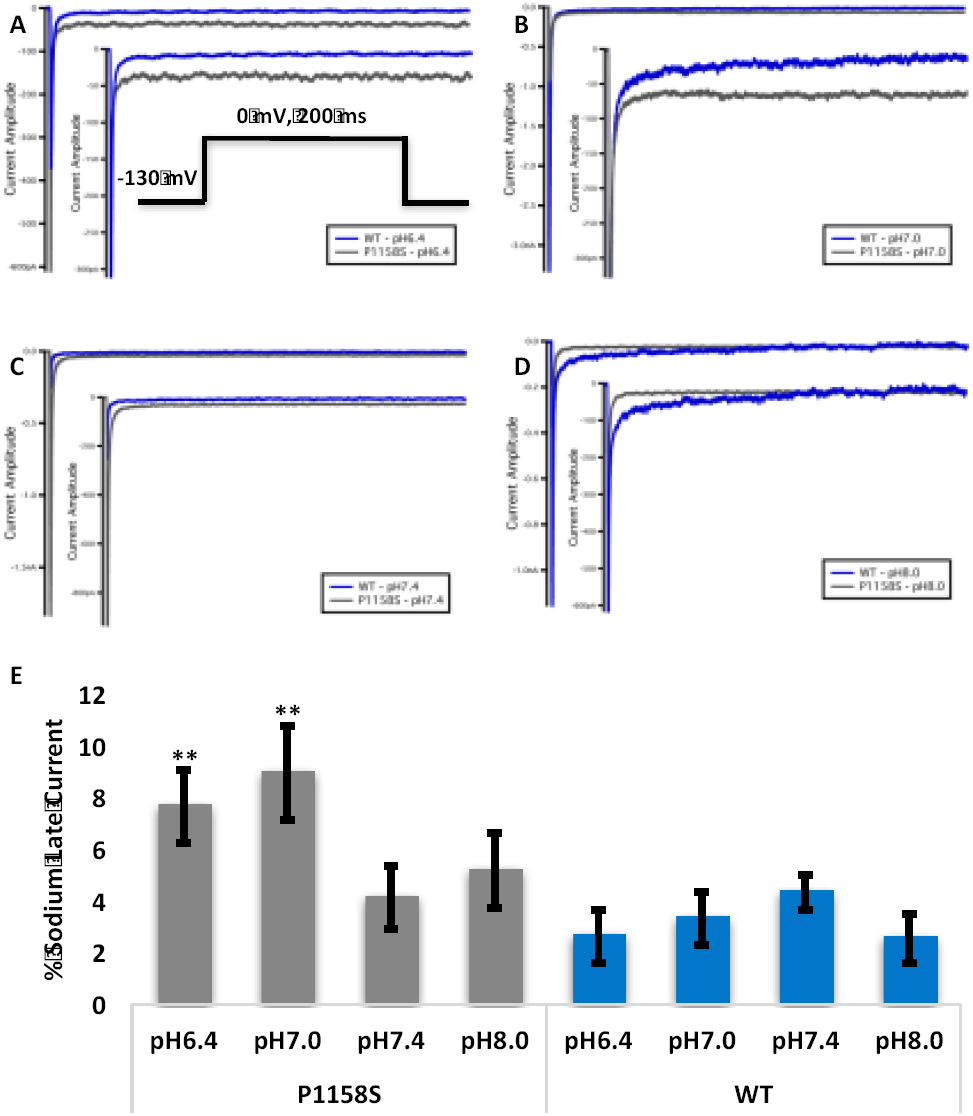
Late sodium current. **(A-D)** Representative normalized current traces of late currents in WT and P1158S at extracellular pH between 6.4 and 8.0. The *inset* in panel **(A)** shows the pulse protocol used to measure late sodium current. **(E)** Shows average late sodium current as a percentage of peak sodium current for WT and P1158S at extracellular pH between 6.4 and 8.0. ** indicate p < 0.01.

To investigate the effects of pH on inactivation in WT and P1158S, we compared use-dependent current reduction at all four pH points. The maximal voluntary contraction frequencies in the soleus muscle, biceps branchii, and deltoid muscle are: ∽11, ∽23, and ∽29 Hz, respectively (38–41). We chose a 45 Hz pulse stimulation frequency to emulate physical activity. Our results suggest that, although WT current decay does not significantly differ with changes in extracellular pH (p > 0.05), P1158S shows a significant acceleration of use-dependent inactivation when extracellular pH is reduced from pH8.0 to 6.4 (p = 0.0142) (**Fig. 6**) (**Table S6**).

**Fig. 6.**
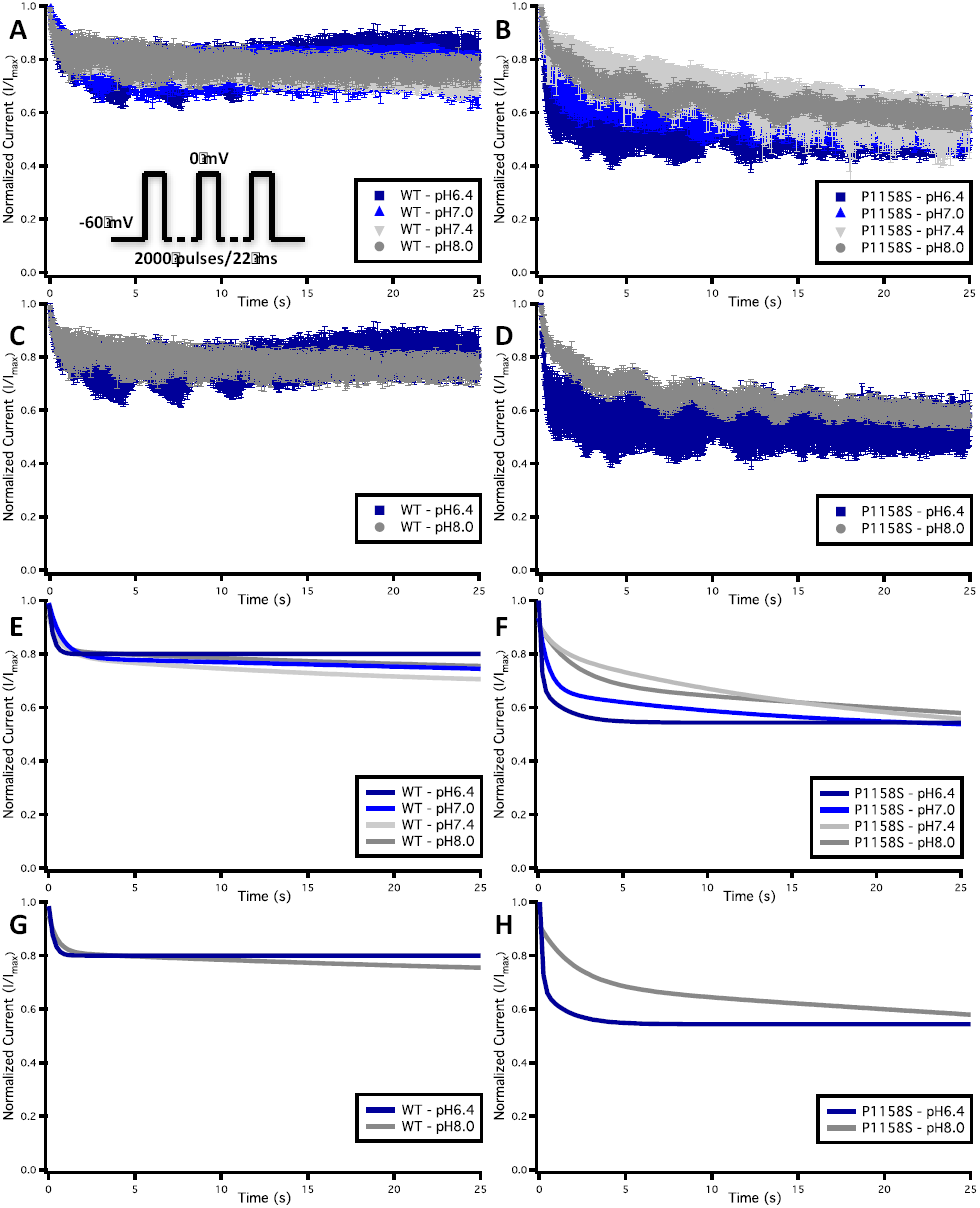
Use-dependent inactivation. **(A-D)** Show normalized current decay plotted as a function of time. The *inset* in panel **(A)** shows the pulse protocol used for these measurements. **(E-H)** Show normalized current decay fitted with an exponential curve.

During physical activity, accumulation of lactate and CO_2_ reduces extracellular pH because of acid efflux from muscle cells (42). These changes in blood acidity play a vital role in blood flow homeostasis (42). However, this acidosis begins on the intracellular side of muscle cells. To test whether the P1158S responses to pH changes are due to protonation on the intracellular side of the channel, we performed whole-cell voltage-clamp experiments in which the pipette solution pH was lowered to 6.4, while buffering the extracellular bath pH at 7.4. The results of these experiments were compared to the experiments with both intra-and extracellular solutions at pH7.4. We found that intracellular acidosis does not significantly alter activation, steady-state fast inactivation, current density, or late current levels in P1158S (p > 0.05) (**Fig. S2**) (**Table S7**). Thus, the observed pH-dependence in the previous experiments are due to extracellular modifications.

### P1158S does not Alter Channel Trafficking or Expression

Our patch-clamp results indicated the average current density of P1158S mutants was significantly greater than that of WT at physiological pH (p = 0.0308) (**Table S2**). Therefore, we sought to determine whether there are any differences in the trafficking and expression between WT and P1158S. We performed immunocytochemistry and used confocal microscopy to visualize and quantify channel distribution and, thus, channel trafficking (**Fig. 7A-C**). Pixel intensity grey value was used as a function of the length of cells (distance) from the middle confocal plane image. For a typical CHO cell with a length of about 14 μm, both WT and P1158S were localized within the 0 to 4 μm and 9 to 14 μm ranges. These two ranges correspond to the outer cellular edges, indicating that most of the channels seem to be concentrated in the proximity of cell membranes (**Fig. 7D-E**). To quantify expression, we measured the average pixel intensity across all confocal planes per cell area, for all cells in all conditions. We found no significant difference between WT and P1158S channel expression (p > 0.05) (**Fig. 7F**).

**Fig. 7.**
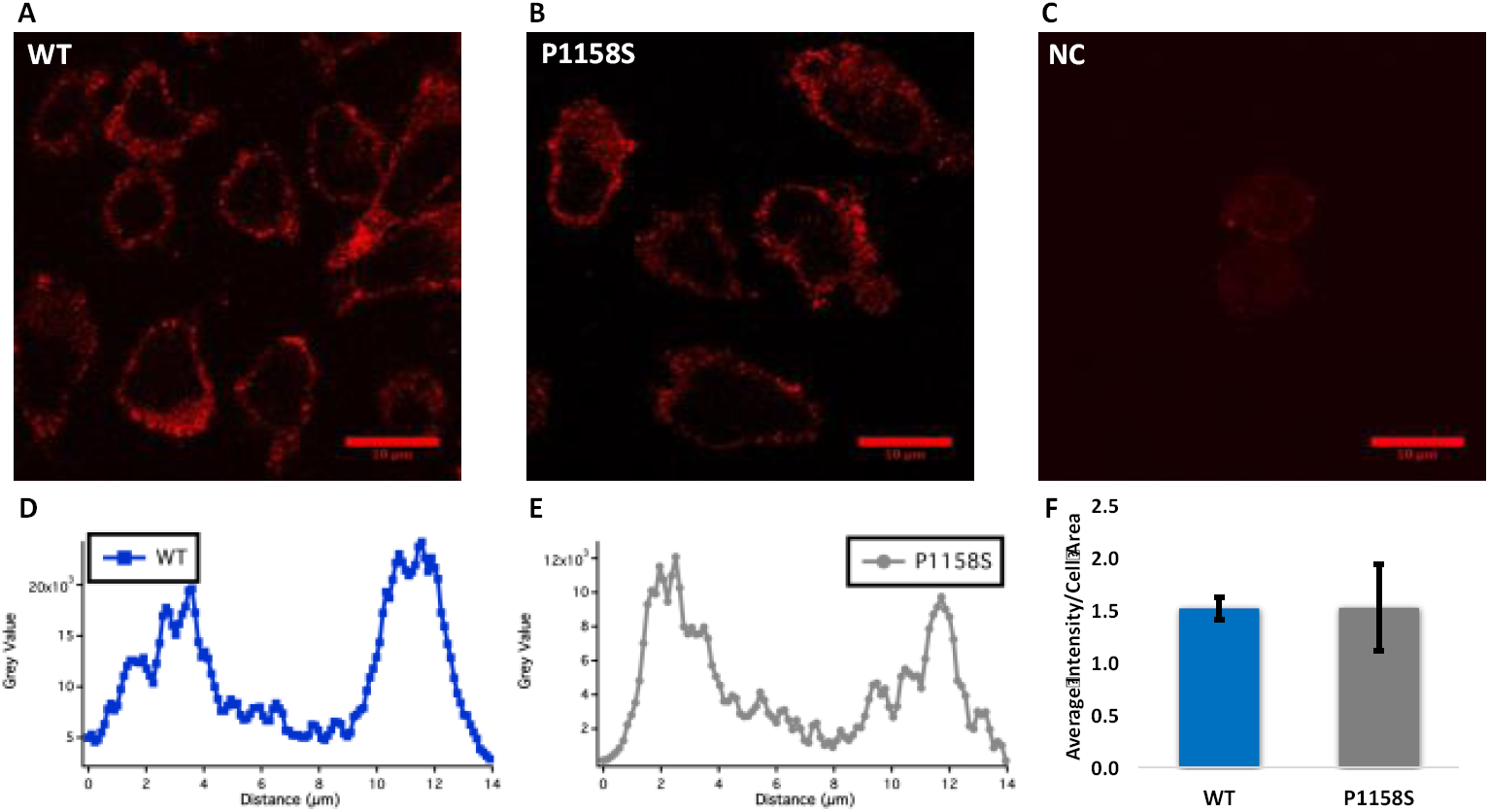
Immunocytochemistry measurements of channel trafficking and expression. **(A-C)** Representative images of WT, P1158S, and negative control cells taken from the middle confocal plane. **(D-E)** Show the distribution of channels, measured by using pixel grey value, across the cell. Both WT and P1158S channels are localized around the edges of cells, in proximity to cell membrane (distances 0-4 μm, and 9-14 μm). **(F)** Average pixel intensity across all confocal images per cell area (n = 3 per condition).

### pH Alterations Can Push P1158S into Periodic Paralysis or Myotonia

We used an action potential model to simulate how the pH-dependent electrophysiological shifts in P1158S may affect phenotype (31). We ran the simulations at a stimulus of 50 μA/cm^2^. A single action potential at physiological pH7.4 required a shorter stimulus duration in P1158S (1.2 ms) than WT (2.0 ms) (**Fig. 8A**). The repolarization of P1158S was slightly prolonged compared to WT due to the presence of larger late sodium current. This late current also led to an increase in the T-Tubule potassium concentrations in P1158S (**Fig. 8B**). The long duration simulation pulse started at 50 ms and stopped at 350 ms. During this pulse, the WT channels activated at 50 ms. WT channels fired only a single action potential. The channels remained inactivated until the stimulus was removed at 350 ms (**Fig. 8C**). However, the action potential morphology changed as a function of pH in P1158S. At the beginning of the stimulus, in pH8.0 models, P1158S mutants displayed an action potential spike, followed by a period during which the membrane potential remained depolarized at −30 mV even after the stimulus was removed (**Fig. 8D**). This inability to repolarize holds the sodium channels in an inactivated state, and is consistent with the periodic paralysis phenotype (36). As with pH8.0, P1158S also displayed a periodic paralysis-like action potential at pH7.4 (**Fig. 8D**). At pH7.0, P1158S showed a continuous train of action potentials for most of the stimulus duration. After the stimulus was removed, there was an after-discharge consistent with myotonic action potentials, which then degenerated into periodic paralysis (36, 43) (**Fig. 8D**). At pH6.4, the P1158S-mutants showed a continuous train of action potentials for the entire stimulation period. After the stimulus was removed, P1158S displayed a progressive repolarization of the membrane potential during a myotonic burst (36, 43) (**Fig. 8D**). Overall, during extreme acidosis, at pH6.4, P1158S only displayed the myotonic phenotype (**Fig. 8E**); however, at the less extreme pH of 7.0, P1158S displayed both myotonia and periodic paralysis (**Fig. 8F**). At pH7.4 and pH8.0, P1158S only showed periodic paralysis (**Fig. 8G-H**).

**Fig. 8.**
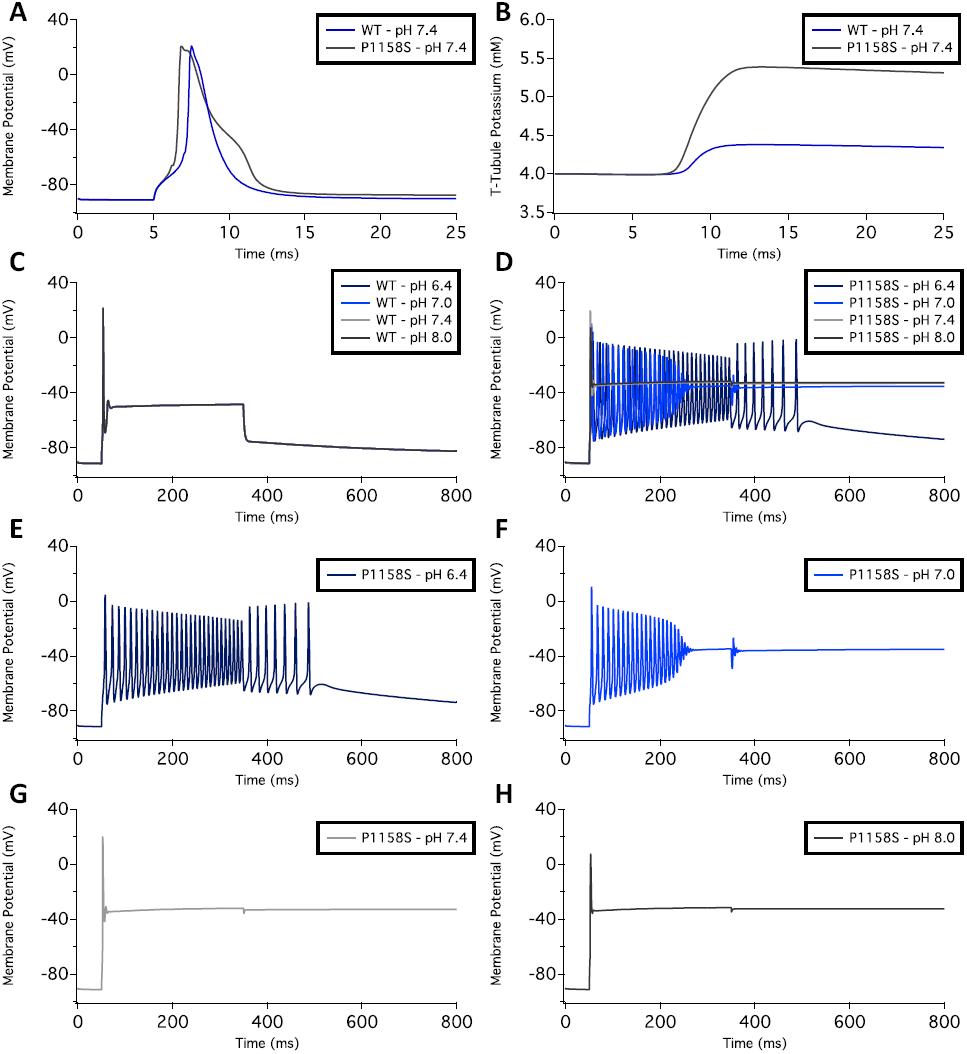
Skeletal muscle action potential modeling. **(A)** Shows single action potential simulations of WT and P1158S channels at the physiological pH of 7.4. The stimulation pulse of 50 μA/cm2 was applied for 2.0 ms in WT and 1.2 ms for P1158S. **(B)** Shows the T-Tubule potassium concentration in WT and P1158S. The potassium concentration is proportional to amount of sodium late currents. **(C-D)** Show long stimulation pulse from 50-350 ms in WT **(C)**, and P1158S **(D)** at all four pH points. **(E-H)** Show P1158S at pH6.4 **(E)**, pH7.0 **(F)**, pH7.4 **(G)**, and pH8.0 **(H)**.

## DISCUSSION

Periodic paralysis and myotonia represent two life-limiting conditions. Interestingly, they fall at the opposite extremes of the disease-spectrum caused by Nav1.4-related channelopathies. P1158S is a single missense mutant that can cause both conditions. We sought to understand how replacing one amino acid might cause both myotonia and hypoPP. Our investigations began with a quest to find common ground between myotonia and periodic paralysis. Because exercise is a common trigger for both conditions, we focused on physiological changes caused by exercise. These changes include variations in cytosolic calcium levels, blood sugar, pH, body temperature, etc. Previous studies characterized P1158S with respect to its temperature sensitivity. A study by Sugiura et al. (2003) suggested the voltage-dependence of activation in P1158S hyperpolarizes upon cooling (21). Furthermore, P1158S can also disrupt slow inactivation at cold temperatures (44).

P1158 is fully-conserved in all mammalian Nav subtypes, and is also in the eukaryotic cockroach sodium channel (NavPaS) (45). The mutation of this vital proline to leucine (P1308L) in Nav1.7 has been implicated in causing inherited erythromelalgia (IEM). IEM is characterized by intermittent burning pain and skin redness in hands and feet. This condition is usually triggered by warmth or exercise (46). Therefore, P1308L in the neuronal Nav1.7 shares some of the triggers with P1158S in skeletal muscles.

We examined whether exercise-induced acidosis can modulate this P1158S. We previously found that Nav1.4 is relatively unaffected by extracellular acidosis compared to Nav1.2 and Nav1.5 (25). The pH-independent current was suggested to protect skeletal muscle activity during exercise-induced acidosis (26). Studies using positron emission tomography indicate that, during exercise, the cardiac glucose consumption is not significantly increased compared to skeletal muscles. This is due to the heart’s ability to use lactate as an alternative source of fuel (47). Therefore, in contrast to skeletal muscles, cardiac cells have a greater ability to respond to acidosis.

We hypothesized that the mutation of P1158 to a serine would change the mobility of DIII. To test this hypothesis, we took a bottom-up approach. We started with a homology model to understand where the conserved P1158 is located on a molecular level. The model also enabled us to determine that there is a hinge-angle difference in the S4-S5 linker with serine instead of proline. Our next step was to find out how much this angle difference would affect the electrophysiology of Nav1.4. The patch-clamp experiments indicated that P1158S causes Nav1.4 to become a pH-sensitive sodium channel, with similar pH-sensitivity to Nav1.2 and 1.5 (25). During extracellular acidosis, the voltage-dependence of activation and steady-state fast inactivation depolarized, current density decreased, and late currents were larger.

Previous studies on Nav1.5-mutants, S1787N and S1103Y, have shown that intracellular acidosis can modulate channel gating leading to arrhythmias (48, 49). To determine whether intracellular pH changes can also modify the gating characteristics of P1158S in Nav1.4, we repeated experiments under acidic intracellular conditions. We found intracellular acidosis to have minimal effects. This suggests the pH-dependence in P1158S is due to the unmasking of protonateable extracellular residues by the mutation.

Changes in the sodium channel expression in NG108-15 cells can alter current density, leading to downstream changes in action potential generation (50). We observed a difference in current density between WT and P1158S, raising the possibility that the mutant might alter channel trafficking and expression. Our confocal microscopy results eliminated this possibility, leaving the mutant effect on channel gating as the most parsimonious explanation for differences in current density.

Lastly, to understand how the pH-dependent changes in P1158S can cause the described phenotypes, we ran action potential simulations. These simulations predict that during alkalosis, such as while hyperventilating (51), P1158S can cause periodic paralysis. However, in the presence of a long stimulus, extreme acidosis can trigger myotonia in P1158S. This is interesting because it is suggested that a pH decrease in muscle cells could alleviate paralytic attacks in some patients (52). Furthermore, P1158S requires a shorter stimulation period to activate compared to WT-Nav1.4. The simulated action potential patterns in P1158S are consistent with the phenotypes associated with periodic paralysis-and myotonia-specific mutants (36, 43).

The molecular mechanism of Nav1.5-proton interactions suggest two mechanisms of proton block. The first mechanism involves the selectivity filter. There are four residues in the P-loop that compose the selectivity filter (DEKA), which along with another four carboxylate residues (EEDD) cause sodium permeation (53–55). These residues are fully-conserved across the sodium channel superfamily. The carboxylates of these residues can get protonated during acidosis. The second mechanism involves the protonation of C373. The pH-insensitivity of Nav1.4 has been attributed to the absence of C373, and the presence of the homologous Y401 in its place (24, 26).

In addition to the two modes of proton block, a third mode of action exists where protons interact with channel voltage sensors (56). We propose that the interaction of protons with the channel voltage-sensors can occur in Nav1.4. Based on the sliding-helix model (32), our results suggest that the mechanism through which P1158S imparts pH-sensitivity in Nav1.4 involves a change in mobility of the DIII S4-S5 linker. Indeed, gating current studies indicate that in Nav1.4, DIII activates prior to DI and DII, suggesting that changes in this linker’s structure could have downstream effects on the mobility of the rest of the channel (28). As a result, the outer vestibule residues that are normally hidden from protons during acidosis, become exposed, and thus mediate pH-sensitivity.

Cations at the extracellular side of the sodium channel VSD modulate gating charges and fast inactivation, hence the presence of protons impedes the immobilization of gating charges (56). Previous studies on Nav1.5 indicate a destabilized fast inactivation at low pH. This suggests that protons disrupt charge immobilization through a direct interaction with the extracellular carboxylates of DIII and DIV VSDs (24, 56). Interestingly, unlike cardiac tissue, the skeletal muscle pH changes more frequently, which may have contributed to the evolution of the pH-insensitive current in Nav1.4 (26, 27, 40). We suggest that the P1158S mutant likely exposes the carboxylates of the DIII VSD, pushing the Nav1.4 gating properties towards Nav1.5.

In conclusion, we characterized a naturally occurring mutation that increases pH-sensitivity in Nav1.4. From a clinical perspective, we identified pH as a potential trigger for periodic paralysis and myotonia in patients with P1158S. Any activity that alters the blood pH balance in P1158S patients could trigger the mutant phenotypes.

## CONFLICT OF INTEREST

None. The authors declare that this research was conducted in the absence of competing interests.

## AUTHOR AND CONTRIBUTORS

M-RG, MA, and CHP collected, assembled, analyzed, and interpreted the data. M-RG wrote the first draft of the manuscript. PCR conceived the experiments and revised the manuscript critically for important intellectual content.

## ACKNOWLEDGMENTS

This work was supported by a grant from Natural Science and Engineering Research Council of Canada and the Canadian Foundation for Innovation to PCR.

Thanks to Dr. Stephen C. Cannon for providing us with the P1158S cDNA, guidance and assistance in action potential modeling. Thanks to Dr. Cynthia Gershome for assistance with ICC.

